# *In vitro* and *in silico* identification of the mechanism of interaction of antimalarial drug – artemisinin with human serum albumin and genomic DNA

**DOI:** 10.1101/519710

**Authors:** Siranush Ginosyan, Hovakim Grabski, Susanna Tiratsuyan

## Abstract

Artemisinins are secondary metabolites of the medicinal plant *Artemisia annua*, which has been traditionally used in Chinese medicine. Artemisinins have anti-inflammatory, anticarcinogenic, immunomodulatory, antimicrobial, anthelmintic, antiviral, antioxidant, and other properties. Our preliminary reverse virtual screening demonstrated that the ligand-binding domain of the human glucocorticoid receptor (LBD of hGR) is the optimal target for artemisinin. At the same time, the binding sites for artemisinin with the ligand-binding domain of the human glucocorticoid receptor coincide with those of dexamethasone. However, the pharmacokinetics, pharmacodynamics, and exact molecular targets and mechanisms of action of artemisinin are not well known. In this work, the interaction of artemisinin with human serum albumin (HSA) was studied both *in vitro* and *in silico*. The results indicate that artemisinin leads to a decrease in optical absorption and quenching of fluorescence by a static mechanism, which is similar to the effect of dexamethasone. Artemisinin interacts with Drug site I on HSA and forms a hydrogen bond with arginine 218. Retardation of the genomic DNA of sarcoma S-180 cells show that artemisinin does not interact directly with DNA. On the basis of the obtained data, we proposed a hypothetical scheme of the mechanisms of action of artemisinin.

**Highlights:**

- Artemisinin quenches the fluorescence of HSA by a static mechanism.
- Artemisinin quenches fluorescence of tryptophan.
- The optimized HSA structure was obtained through molecular dynamics simulations.
- Artemisinin binds with HSA in Drug site I and forms a hydrogen bond with Arg218.
- Dexamethasone binds with HSA in Drug site I and forms hydrogen bonds with Arg218, Arg222 and Va1343.
- A hypothetical scheme of the mechanism of action of Artemisinin was proposed.

**Graphical Abstract:** 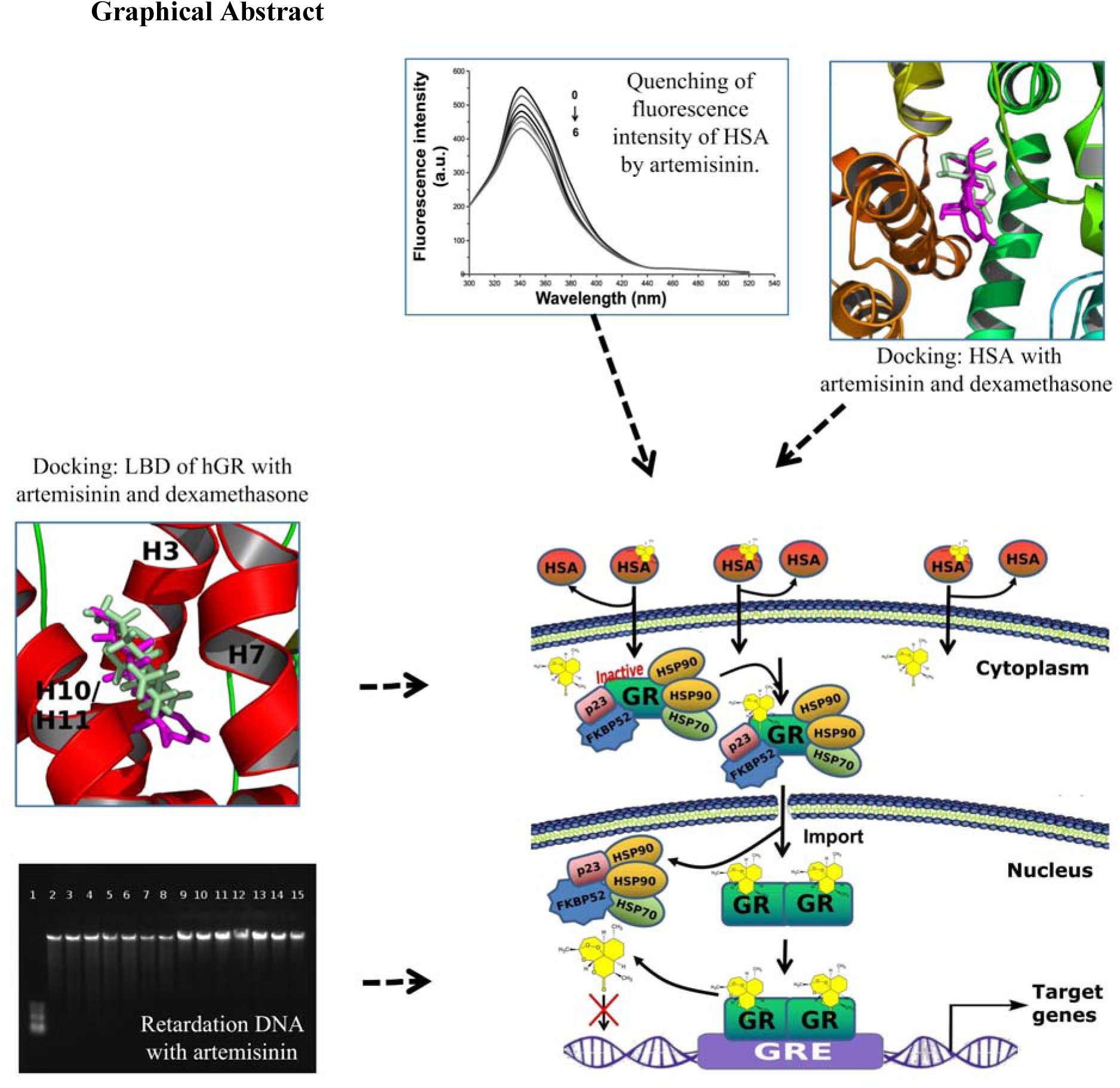

## 1. Introduction

Artemisinins (ART) belong to the family of sesquiterpene trioxane lactones, the secondary metabolites of the herb plant *Artemisia annua* with potent antimalarial, antiviral, antifungal, antitumor and anti-inflammatory properties and used in traditional medicine [1, 2]. Artemisinin (C_15_H_22_O_5_) [3] contains an endoperoxide bridge which is necessary for its biological activity (C-ring) [4]. Artemisinins exhibit various activities, such as antioxidant, anti-inflammatory, immunomodulatory, antimicrobial, antihelminthic, antiviral and other [5-8]. In 2015, the Nobel Prize was received for using it as an antimalarial drug [9]. Artemisinin and its derivatives exhibit sustained inhibitory effect on fungi; *in vitro* and *in vivo* cytotoxic activity against cancer cells, such as leukemia, melanoma, breast, ovarian, prostate gland, kidney cancer cell lines and plant tumors [10]. Nowadays ART is considered as a therapeutic alternative in the treatment of rapidly progressive and highly aggressive forms of cancer without the risk of acquiring resistance [11]. Artemisinin can also reduce glucose levels and act against diabetes mellitus [12].

It has been shown that ART has antibacterial properties against different bacterial species [13]. It has been also shown that derivatives of ART such as artesunate can enhance the effect of antibiotic methicillin against resistant *Staphylococcus aureus* and beta-lactams against *E.coli*. This is achieved through the inhibition of multidrug-resistance efflux pumps (MDR) AcrAB-TolC, which leads to the suppression of pro-inflammatory cytokines TNF-α, release of IL-6 and increasing the accumulation of antibiotics in bacteria [14]. One of the possible mechanisms of the antimalarial activity of ART and its derivatives could be the accumulation and release of free radicals due to the destruction of the endoperoxide bridge by the intracellular heme-bound iron (Fe). [15]. However, up to now, the pharmacokinetics, pharmacodynamics, specific molecular targets and mechanisms of action of the ART family compounds have not been well studied [4-6, 8].

Our preliminary reverse virtual screening demonstrated that the optimal target for ART is the ligand binding domain of the human glucocorticoid receptor (LBD of hGR) [16]. It is a constitutively expressed transcription factor controlling a variety of different gene networks [17, 18]. Ketosteroid receptors, including hGR, are activated by endogenous hormone cortisol or exogenous glucocorticoids (GC), such as dexamethasone (DEXA) and play a vital role in the maintenance of metabolic and homeostatic regulation [19, 20], in regulation of cell-cell communication required for the coordination of development, growth, inflammation and immunity [21, 22]. Therefore, GC and their synthetic derivatives are one of the most used drugs in the treatment of asthma, rheumatoid arthritis, autoimmune diseases, leukemia and others due to the strong immunosuppressive, anti-inflammatory, and apoptotic activity. But the use of steroids can lead to serious adverse effects [23]. Therefore, novel therapeutic agents are researched that can replace steroids and be used as anti-inflammatory drugs and not cause side effects. These compounds can be selective GR agonists or selective GR modulators [20]. Thus development of new drugs requires an understanding of the molecular mechanisms of hGR action.

Most xenogeny factors, drugs such as native and synthetic GC [24] and ART [25] reach the target tissues by binding to human serum albumin (HSA), which is involved in the absorption, distribution, metabolism and excretion of drugs. Human serum albumin has significant effect on the pharmacokinetics and pharmacodynamics of many drugs [26]. Suction of ART after ingestion lasts 0.5-2 hours, reaching a maximum concentration in the plasma in the interval of 1-3 hours after administration. Enzymes of cytochrome P-450 CYP3A4 and CYP2A6 are involved in the metabolism of ART and its derivatives. They are also involved in the conversion of ARTs’ derivatives into dihydroartemisinin [27].

Human serum albumin is a single-chain, non-glycosylated polypeptide that contains 585 amino acids with a molecular weight of 66.500 Da. It forms a heart-shaped structure consisting of three homologous α-helical domains (I, II and III), each of which is subdivided into two subdomains (A and B) [28]. Analysis of the crystal structure showed that there are two large and structurally selective binding sites - the Sudlow I site (Drug site I) and the Sudlow II site (Drug site II), which are located respectively in the IIA and IIIA subdomain [29].

In this paper various methods are used to study the interaction between small molecules and HSA, such as UV spectroscopy, fluorescence spectroscopy [26, 30] and molecular docking [31]. The objectives of this work were *in vitro* and *in silico* determination of the molecular mechanisms of the interaction and binding of ART with HSA, genomic DNA and comparative analysis with DEXA.

## 2. Materials and Methods

### 2.1. Reagents and Solutions

The standard solutions for ART, HSA and tryptophan (TRP) were prepared using the following: ART (≥98%) of the firms TRC (Toronto, Canada); HSA, free from fatty acids (<0.005%) was purchased from Sigma Chemical Co, USA; TRP and tris-(hydroxymethyl) aminomethane (Tris) (≥99%) from Sigma Chemical Co, USA. Tris-HCl buffer solution (0.050 M, pH = 7.40) was adjusted to pH = 7.40 by adding 36% HCl. The initial solution of ART (1.0×10^−2^ M) was prepared in ethanol. The initial solution of HSA (2.5×10^−4^M) was prepared in Tris-HCl buffer solution (pH = 7.40). The HSA concentration (0.4 mg/ml) was determined by spectroscopic method at a wavelength of λ = 280 nm using a molar absorption coefficient of ε = 36.600 M^-1^cm^-1^. All initial solutions were kept in dark environment at 4 °C. Distilled water was used during the experiments.

### 2.2. Spectroscopic methods

The UV spectra of all HSA solutions in the absence/presence of ART under physiological conditions (pH=7.4) were recorded on Unicam UV-1601 spectrophotometer (Shimadzu, Japan) with 10-millimeter quartz cells in the range from 200 to 350 nm at room temperature. Standard solutions of HSA and ART were used as references. The fluorescent spectrum of HSA solutions in the absence/presence of ART under physiological conditions (pH = 7.4) were recorded on a Varian Cary Eclipse Fluorescence Spectrophotometer with 10 mm quartz cells with a slot width is 10/10 nm. Fluorescence emission spectra were recorded in the range from 300 to 500 nm with an interval of 0.2 nm at an excitation wavelength λ_ex_ = 280 nm at three different temperatures (298, 304 and 310 K). The width of the spectral excitation and emission bands was set to 5 nm. The temperature of the sample was maintained by recycling water throughout the experiment.

Order statistics were used to analyze the obtained data from spectrophotometry. For normally distributed data, methods of parametric statistics were used, which includes parametric statistics and unpaired Student’s t-test. The calculations were performed using Microsoft Office Excel 2007 program. A statistically significant value was taken as p<0.05. Reproducibility was ensured by performing 4-6 runs with 2-3 series of each runs.

### 2.3. DNA Retardation

Artemisinin was dissolved in 10 ml of dimethyl sulfoxide (DMSO) to a final concentration of 1, 5, 10, 25, 50 and 100 μM. Work with line of mice with S-180 sarcoma cells was conducted in the Laboratory of Toxinology and Molecular Systematics of L. A. Orbeli Institute of Physiology NAS RA. Genomic DNA was isolated from a small volume of cells of the S-180 sarcoma cancer line using the peqGOLD MicroSpin Tissue DNA Kit (PeqLab Biotechinologie GmbH Erlangen, Germany). DNA integrity was evaluated in comparison with the molecular weight standard by measuring the electrophoretic mobility of genomic DNA in a 0.8% agarose gel [32].

### 2.3. Molecular modeling

#### 2.3.1. Molecular dynamics (MD) simulations and cluster analysis of HSA

The 3D structure of HSA with complexed with myristic acid and the R-(+) enantiomer of warfarin was taken from the RCSB Protein Data Bank [33] (PDB ID: 1H9Z). The crystal structure of HSA has missing residues and atoms, thus it was necessary to fix it. The addition of missing amino acids was made using the Modeller software package [34]. Molecular dynamics simulations were performed using the GROMACS package [35]. The Amber99SB-ILDN force field [36] and the TIP3P [37] water model were used. Virtual sites were used for increasing the time step to 4 fs. Later energy minimization was performed and MD simulations were carried out in explicit water at a constant temperature (300 K). The charge of the system was neutralized by randomly replacing water molecules with Na and Cl ions. Periodic boundary conditions were applied to the system. Particle-Mesh Ewald (PME) was used for electrostatics with a cut-off of 1.0 Å. The simulation was run 4 times (400 ns each) which resulted a total of 1.6 μs. To obtain the most common HSA conformation, cluster analysis was performed using the Gromos RMSD-based method [38] and the criteria were based on the work λof Offutt et al. [39]. The molecular dynamics simulations were performed on the HPC of Lomonosov Moscow State University [40].

#### 2.3.2. Ligand preparation

3D structures of ART [CID: 68827] and DEXA [CID: 5743] were acquired from PubChem database [41]. Topology of the ligands for the MD simulations was generated using acpype [42], which is compatible with the General Amber Force Field [43]

#### 2.3.3. Blind Docking with multiple programs

Docking experiments of ART and DEXA with HSA were performed using Autodock Vina [44], rDock [45], Ledock [31], FlexAID [46]. The first three software packages were chosen based on their accuracy [47], while Flexaid for its stability against structural variability [46].

The whole protein conformational space was searched, using grid box dimensions 94×54×92 Å for Autodock Vina. Following exhaustiveness values were tested in this study: 8, 16, 32, 64, 128, 256, 512, 1024, 2048 and 4096. So a total of 200 conformations were generated using Autodock Vina. The radius was to 50 Å to cover the whole protein surface. The box size of LeDock was also set to cover the whole protein surface with the following values: x_min_ = -10, x_max_ = 11; y_min_ = -20, y_max_ = 21; z_min_ = -42, z_max_ = 43. As for the FlexAID we strongly followed with the parameters of the template as they were tested and are optimal [46]. Simulations for each program were run 10 times with binding modes set to generate 20 conformations. A total 800 conformations were generated for the interaction of ART with HSA.

#### 2.3.4. Analysis of docking results: conformations and trajectories

The center of mass (COM) coordinates were extracted and then the dimensions of the data were reduced using Principal Component analysis [48]. Later cluster analysis was performed on the COM data using the Density-based spatial clustering of applications with noise (DBSCAN) algorithm [49], which is a well-known data clustering algorithm that is commonly used in data mining and machine learning. Blind docking methodology was based on a previous study [50], which uses pandas [51], scikit-learn [52] and openbabel [53]. After that centroid conformations were extracted from the common clusters for all the programs. Later the centroid conformations were used as the foundation for local docking. Local docking was performed with Autodock Vina [44]. Analysis of the interactions and hydrogen bonds was done using Ligplot^+^ [54]. Visualization of the docking conformations was performed using Pymol [55].

## 3. Results and Discussion

### 3.1. UV-vis spectroscopy

The UV absorption spectra of HSA with different concentrations of ART are presented in Fig 1a.

**Fig. 1.**
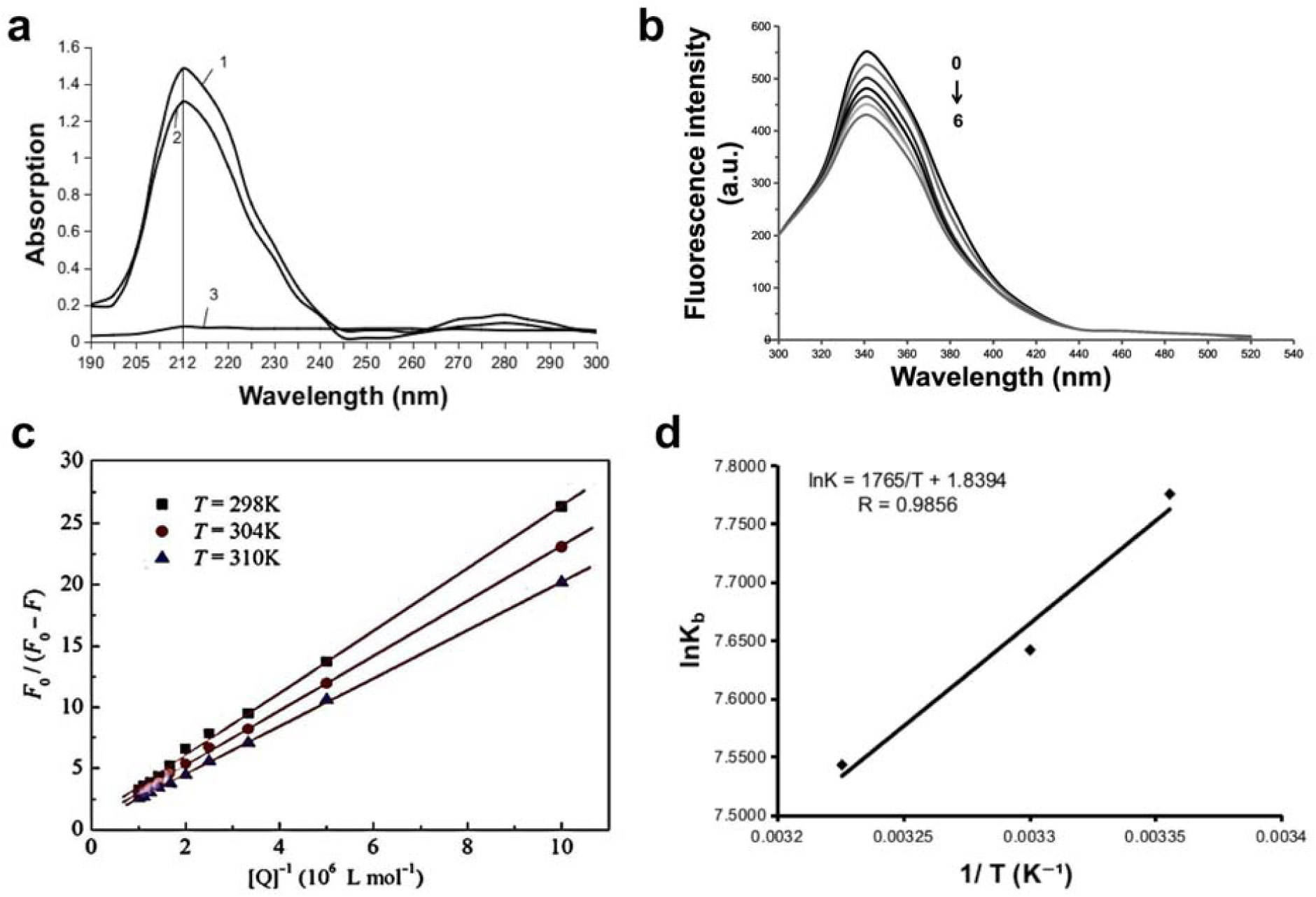
a) UV spectra of HSA (1) 1.5×10^−6^ M (as a control- buffer solution); (2) HSA+ART (12×10^−6^ mol L^−1^ (1:1); (3) ART (2×10^−6^ mol L^−1^), pH 7.4, 298 K. b) Fluorescence emission spectra of the HSA (1.0×10^−6^ M) and ART, λ_ex_ = 289 nm, 298 K. 0-6 - the concentrations of ART 10^−5^×0, 2, 4, 6, 8, 10,12 mol L^−1^,respectively. c) Modified Stern-Volmer plots for the quenching of HSA by ART at 298 K (R=0.9995), 304 K (R=0.9975); 310 K (R=0.9965). d) Van’t Hoff plots for the interaction of HSA with ART, pH 7.4, (Tris–HCl buffer, [HSA] = 1.5×10^−6^ mol L^−1^).

As it can be seen from Fig. 1a, the HSA has two absorption peaks: strong absorbance at about λ = 212 nm and weak at 280 nm. The first band reflects the conformation of HSA and the second - belongs to the π→π* transition of the aromatic amino acids, specifically tryptophan [30]. The intensity of absorption band near 212 nm decreased with a slight red shift of absorption band with the addition of ART. Meanwhile the intensity of band near 280 nm slightly decreased, which indicates a slight change in the conformation of HSA and the content of α-helix due to the formation of ART–HSA complex. This indicates the interaction of ART with HSA. Information on the conformational changes of HSA before/after the addition of ART was obtained by measuring the intensity of fluorescence.

Human serum albumin has fluorescent properties and emits light intensively upon excitation, and three amino acid residues tryptophan, tyrosine and phenylalanine are responsible for the fluorescence [30, 56]. Human serum albumin contains only one Trp214 residue in the large hydrophobic cavity of the subdomain IIA, which is the main fluorescent component and accounts for about 90% of the total protein fluorescence [28, 57]. Human serum albumin generated an apparent fluorescence emission band at 337 nm, which decreased and gradually damped with increasing ART concentration without a significant shift (Fig. 1b). Moreover, ART did not fluoresce even at the maximum concentration. Consequently, such a decrease in intensity may be due to quenching of fluorescence of HSA by ART, as a result of the complex formation. In order to confirm the quenching mechanism, we analysed the fluorescence data at different temperatures with the Stern–Volmer equation [30]:

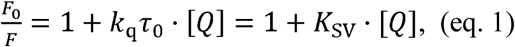

where *F*_0_ и *F* - are the fluorescence intensities of HSA in the absence and in the presence of the quencher (ART). *K*_SV_ – is the Stern-Volmer quenching constant, [*Q*] is the concentration of the ART, *k*_q_ is the quenching rate constant of the biomolecule and τ_0_ is the average lifetime of the molecule in the absence of the ART. A linear *F*_0_*/F* vs. [*Q*] plot indicates that a single type of quenching mechanism is involved, while deviation from the linearity suggests a mixed quenching mechanism [30]. Modified Stern-Volmer plots were used to analyse the mechanism (static or dynamic) of fluorescence quenching. Based on the calculated fluorescence data at three temperatures (298, 304 and 310 K), *F*_0_/*F* graphs are plotted relative to [*Q*] (Fig. 1c):

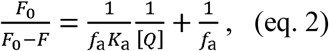

where *K*_a_ is the effective quenching constant for the accessible fluorophores and *f*_a_ is the fraction of accessible fluorophore. The plot of *F*_0_/*F*_0_-*F* vs. 1/[*Q*] yields 1/*f*_a_ as the intercept, and 1/(*f*_a_*K*_a_) as the slope. k_q_ = K_SV_/τ_0_, where τ_0_ - the lifetime of HSA without quencher ART. The lifetime of the biopolymer fluorescence is 10^−8^ sec.

As can be seen in Fig. 1c, within the range of studied concentrations, the results agree with the Stern-Volmer equation, and the graph shows linear relationship. This testifies in favor of the fact that in the presence of ART the quenching of HSA proceeds according to one mechanism and secondly, with increasing temperature, the quenching rate decreases. This indicates the static nature of extinguishing [30].

### 3.2. Binding constants and the number of binding sites

When small molecules bind independently to a set of equivalent sites on a macromolecule, the binding constant (*K*_b_) and the number of binding sites (*n*) can be determined by plotting the double-log regression curve of the fluorescence data with the modified Stern–Volmer equation (eq. 2) [58, 59]:

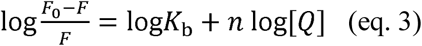

where *F*_0_ and *F* are the fluorescence intensities of the HSA in the absence and in the presence of ART, [*Q*] is the concentration of the ART, *K*_b_ is the binding constant and *n* is the number of binding sites. The *K*_b_ values are obtained from the log [(*F*_0_ -*F*)/*F*] vs. log [*Q*] plot.

Enthalpy change (ΔH), entropy change (ΔS), and Gibbs free energy change (ΔG), were calculated using the van’t Hoff equations which used to characterize the binding force involved in the ART–HSA [30]:

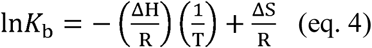

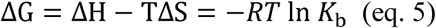

where *K*_b_ is the binding constant at the T=298, 303, 310 K and the gas constant R is 8.314 J/mol·K, which was converted to cal/(mol·K). ΔH and ΔS values were obtained from the slope and intercept of the linear plot ln*K*_b_ versus 1/T based on eqn (4) (Fig. 1d)

The values of ΔH, ΔS and ΔG are listed in Table 1. The negative sign of ΔG indicates that all binding processes are spontaneous. Positive ΔS value is frequently regarded as an evidence for a hydrophobic interaction during drug-protein interaction. Our data corresponds to the study [25].

**Table 1.**
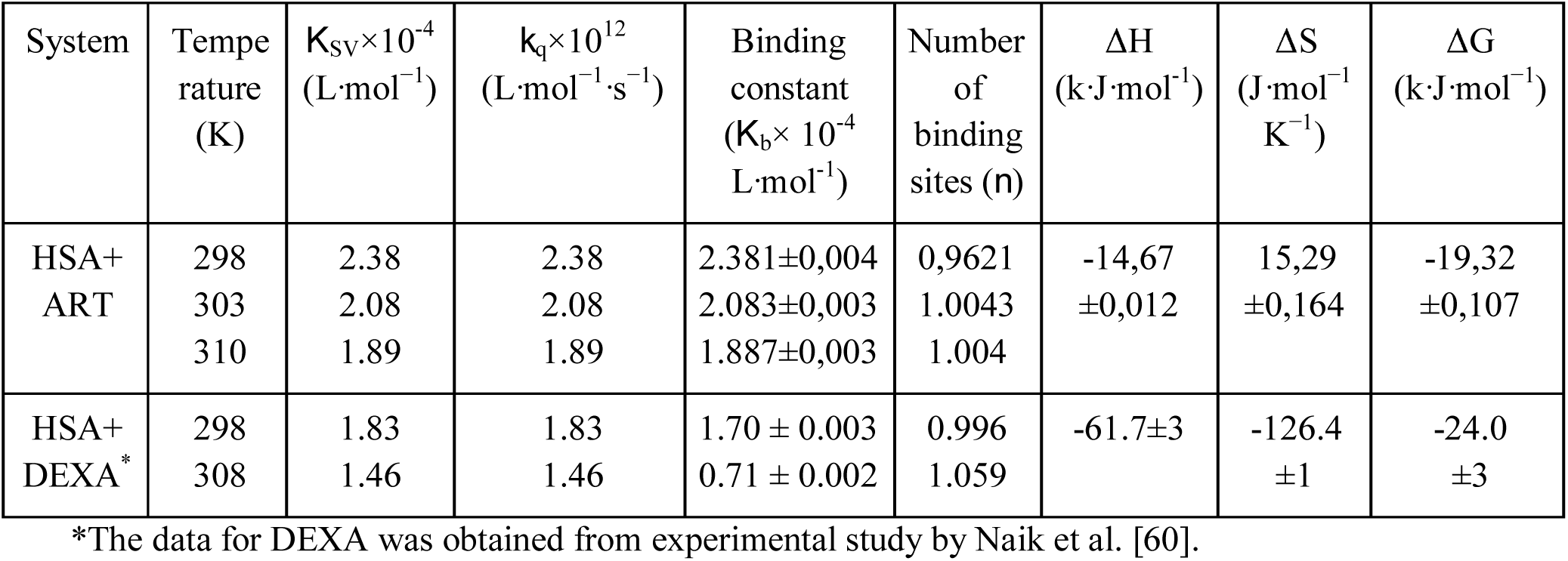
The constants of quenching (*K*_SV_, *K*_q_), binding (*K*_b_) and thermodynamic parameters of HSA-ART and HSA-DEXA interaction at pH 7.40.

### 3.3. Energy Transfer from HSA to ART

The estimation of the interactions, measuring distance between ART (acceptor) and the fluorophore molecule HSA (donor) was carried out based on the Förster’s resonance energy transfer (FRET) theory. The transfer of energy is controlled by the fluorescence quantum yield of the donor (*E*), the relative orientation of the transition dipoles of the donor and acceptor (*k*^2^), overlap integral between the fluorescence spectrum of the donor and absorption spectrum of the acceptor (*J*), and the distance (*r*) between them [61]. The overlap of the fluorescence spectra of HSA and the absorption spectra of ART is shown in Fig. 2a.

**Fig. 2.**
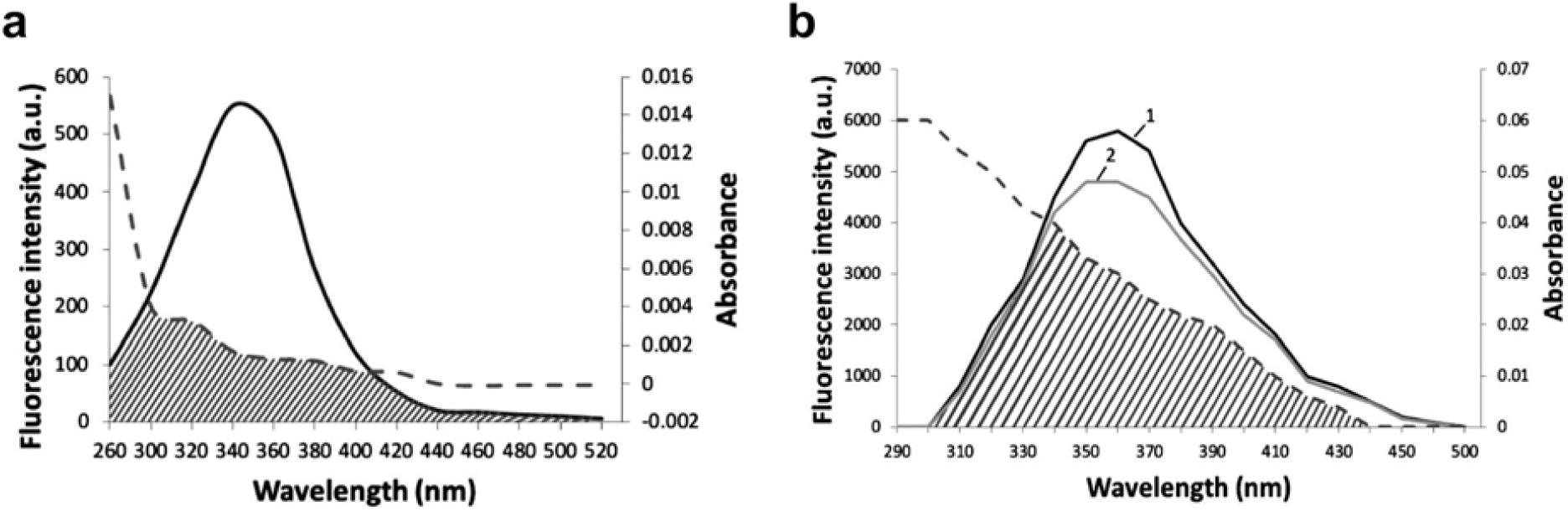
a). Spectral overlap between the fluorescence emission spectrum of HSA (1.0×10^−6^ M) (the straight line) and the absorption spectrum of ART (1.0×10^−6^ M) (the dish line). b) Fluorescence emission spectra of Trp (1) and Trp-ART (2). Spectral overlap between emission spectrum of Trp (the straight line) and the absorption spectrum of ART (the dish line) pH = 7.40, T = 298 K.

The efficiency of energy transfer for a single donor-single acceptor system is expressed by equation [61]:

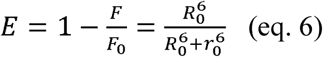

where *F*_0_ and *F* are the fluorescence intensities of HSA in the absence and presence of ART, respectively. *R*_0_ so called the Förster length, which is a function of the mutual orientation of HSA and ART and the degree to which the excitation spectrum of ART overlaps the emission spectrum of HSA. *R*_0_ is the critical distance when the energy transfer efficiency is 50% and its value can be calculated by the following equation [62]:

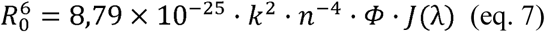

where *k*^2^ is the orientation factor related to the geometry of the donor–acceptor of dipole and *k*_2_ = 2/3 is for the random orientation in the fluid solution, *n* is the refractive index of medium, Φ the fluorescence quantum yield of the donor, *J* is the spectra overlap of the donor emission and the acceptor absorption. *J* is given by equation [62]:

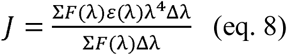

where *F*(λ) is the fluorescence intensity of the fluorescence reagent where λ is the wavelength, ε(λ) is the molar absorbance coefficient of the acceptor at the wavelength of λ. From these relationships, *J, E, R*_0_ and *r* can be calculated.

Energy transfer in the spectra may depend on the distance between the tryptophan residue and ART bound to HSA. The overlap integral calculated from Fig. 2a is 2.5216×10^−15^ cm^3^/mol^−1^. Thus the value of *R*_0_ is 1,98 nm and the value of *r* is 1,89 nm. For DEXA-HSA system *R*_0_ =2.53 nm and *r* = 2.87 nm [60]. The distance between HSA (donor) and ART (acceptor) are much smaller than a criterion value for non-radiative energy transfer phenomenon [63]. This corresponds to the rule 0.5*R*_0_ < *r*_0_ < 1.5*R*_0_ [64], thus suggesting that the energy transfer from HSA to ART occurs with high probability and distance obtained by FRET with high accuracy. The average distance between a donor and acceptor is 2–8 nm, which indicated that the energy transfer from HSA to DEXA as well as ART–HSA occurred with high probability. The larger *R*_0_ value obtained for DEXA–HSA system is indicative of more efficient FRET, which was observed in the DEXA–HSA complex than ART–HSA complex [60].

The effect of ART on the fluorescence intensity of Trp in H_2_O at 293 K is presented in Fig. 2b. Fluorescence of Trp is characterized by maximum emission of λ_em_ = 356 nm. The spectral characteristics of 2D fluorescence spectra of amino acids are recorded at the following conditions: for Trp λ_ex_/λ_em_ = 280/357 nm ranges are given. The UV/vis spectrum of Trp is characterized by a short wavelength band at 220 nm and a long wavelength band at 260–290 nm, which consists of two overlapping transitions. The fluorescence spectrum of Trp in aqueous solution is characterized by a wide structureless emission band with a maximum of 357 nm and width of 60 nm when λ_ex_=280 nm, which decreased and gradually damped with increasing concentration of ART without a significant shift (Fig. 2b). Energy transfer measurements were calculated using equation 6.

For ART-TRP system *J* = 3.6×10^−16^ cm^3^/mol^−1^ and the value of *R*_0_ is 2,07 nm and the value of *r*_0_ is 2,05 nm. The distance between Trp (donor) and ART (acceptor) was much smaller [63] than the criterion value for non-radiative energy transfer phenomenon (1.035 < 2.05 < 3.105).

### 3.4. Interaction of ART with genomic DNA of S-180 cancer cell line

Artemisinin when interacts with GR, can be imported into the nucleus, and it is extremely important to assess the direct interaction of ART with genomic DNA. The study of direct interaction of genomic DNA of S-180 cancer cell line with ART was carried out by measuring electrophoretic mobility of genomic DNA [65] (Fig. 3). Different concentrations and joint incubation times were tested.

**Fig. 3.**
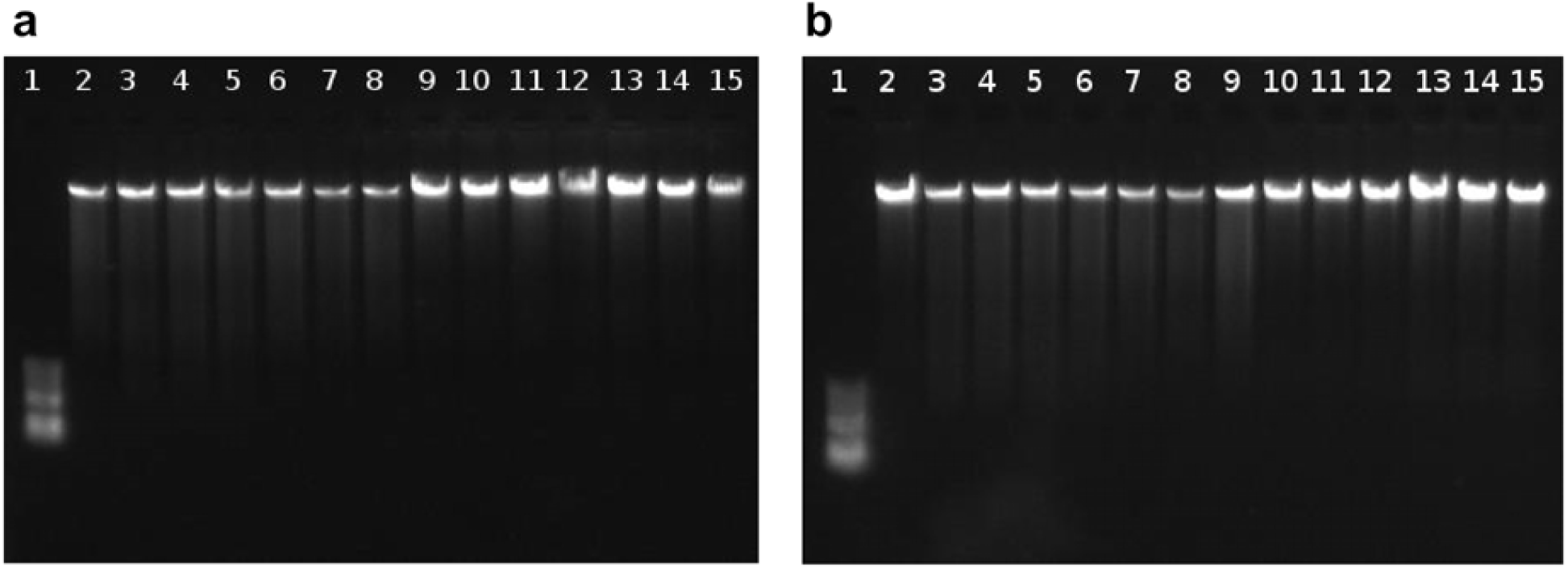
The electrophoretic mobility of the genomic DNA of the S-180 cells during 30 min (a) and 48h (b) of incubation. 1-Ladder; 2,9-DNA; 3-8, 10-15 - DNA with ART: 3,10 - 1 μM; 4,11 - 5 μM; 5,12 - 10 μM; 6,13 - 25 μM; 7,14 - 50 μM; 8,15 - 100 μM ART.

The results of our experiments show that direct interaction of ART with the genomic DNA of S-180 line cells at concentrations ranging from 1-100 μM ART during incubation for 48 hours is not observed. There are conflicting data regarding the direct interaction of ART with genomic DNA. According to one source, direct interaction of ART with DNA is observed, which leads to DNA damage [66]. Our results are consistent with data, which claim that there is no direct interaction between them [4, 10].

### 3.5. Molecular docking of HSA with ART and DEXA

We performed molecular dynamics simulations of HSA after adding the first two missing amino acids. The simulation was run 4 times for 400 ns, which resulted in a total of 1600 ns. Cluster analysis of the simulations was performed using the Gromos RMSD-based method. The Gromos method developed by Daura et al. [38] based on the calculation for each point the number of other points for which the RMSD is less than a given cutoff. In our case, the cutoff value was chosen 0.26, according to the 3 criteria described in [39]. As a result, we obtained 36 clusters, where the first cluster was 43.25% of the total number of conformations (Fig. 4). The distribution of clusters over time shows that the first cluster includes conformations encountered in all four simulation runs (Fig. S1). From which we can conclude that this HSA conformations is not random.

**Fig. 4.**
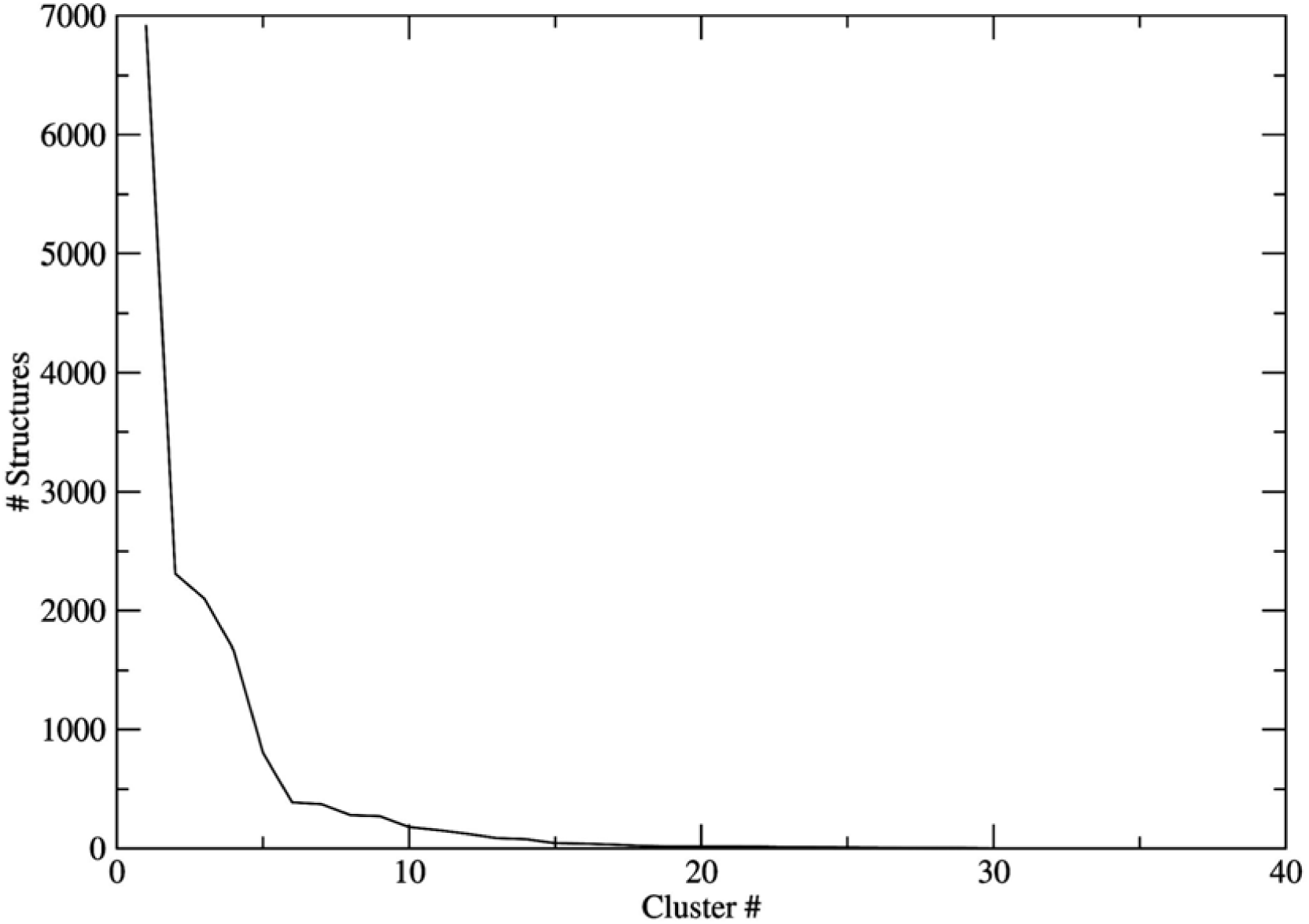
Distribution of HSA conformations.

For further research we used the centroid structure of the first cluster. To study the mechanism of interaction of ART interaction with the centroid conformation of the first HSA cluster, docking simulations were performed using four programs: Autodock Vina [44], rDock [45], Ledock [31], FlexAID [46].

In the case of docking, the simulations were run 10 times and the number of binding modes set to 20 for each program. For each program, 200 conformations of the ligand were obtained, and a total of 800 docking conformations were generated. Large amount of conformations were necessary to obtain good sampling and for further cluster analysis. We performed Principal component analysis of the COM coordinates of artemisinin conformations [50] (Fig. 5a). Then cluster analysis was carried out using the DBSCAN algorithm. DBSCAN algorithm has two important parameters: epsilon and minimum number of points. Parameter epsilon is defined as the radius of neighborhood around a certain point x, and parameter minimum number of points as the minimum quantity of neighbors in the limit of the radius [49]. Minimum number of points was set to 5, and optimal epsilon value was determined based on kNN plot which is equal 2.0 (Fig. S2). As a result of cluster analysis 6 binding regions were determined from the 800 conformations of ART (Fig. S3). One (I) of the six clusters amount the most (58.2%). This means that the results of all 4 docking programs in this area coincided and that this site is a potential binding site for ART with HSA. This site corresponds to subdomain IIA domain or Drug site 1, which corresponds to experimental data [25].

**Fig. 5.**
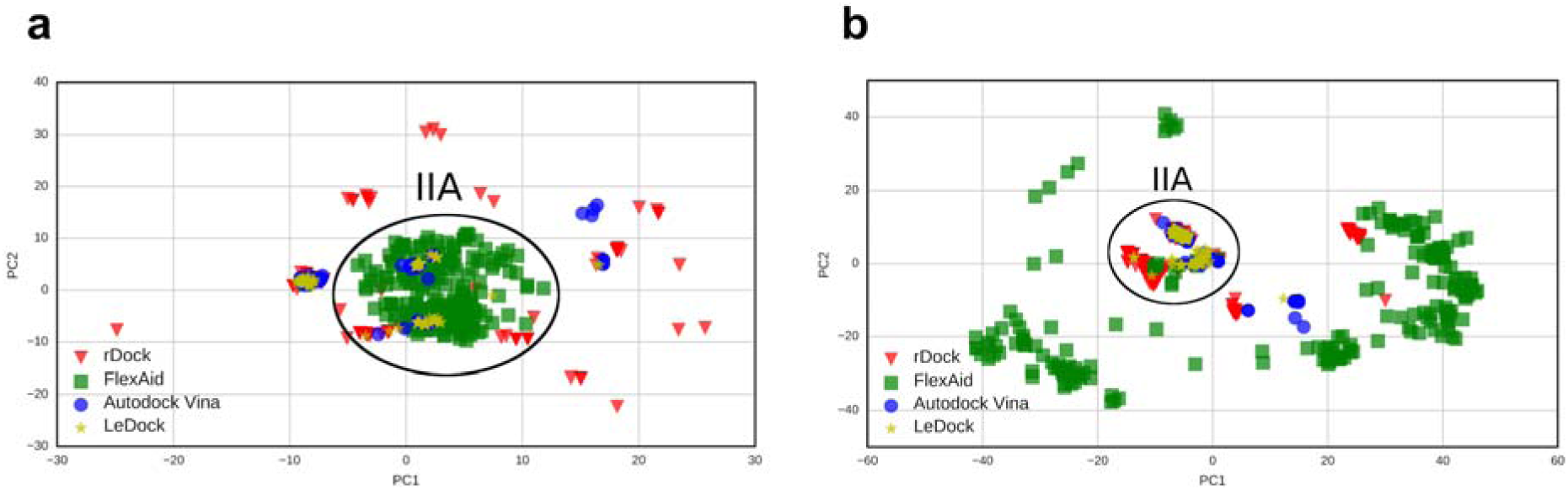
a) PC analysis of the results of multiple docking runs of ART with the centroid conformation of HSA. b) PC analysis of the results of DEXA with multiple docking programs with the centroid conformation of HSA.

After determining the binding site of ART on HSA (IIA), a detailed analysis was performed. Therefore, we conducted local docking using AutoDock Vina [44] (Fig. 6). AutoDock Vina was chosen due to the fact that it had the best scoring power, which is based on empirical free energy and is frequently used in research [47]. The binding energy of the best pose is -8.4 kcal/mol, which is quite high. Analysis of the hydrogen bonds and hydrophobic interactions showed that ART forms many hydrophobic interactions (Gln221, Arg222, Glu292, Asn295 and Val343) and one hydrogen bond with arginine 218, which plays a critical role in the binding of thyroxin [67], warfarin, bilirubin and cholesterol [68]. Artemisinins also form hydrophobic interactions with amino acids from the IIIA domain (Lys444, Pro447 and Asp451). It does not form hydrogen bonds with tryptophan 214, which is confirmed by the results of fluorescence.

**Fig. 6.**
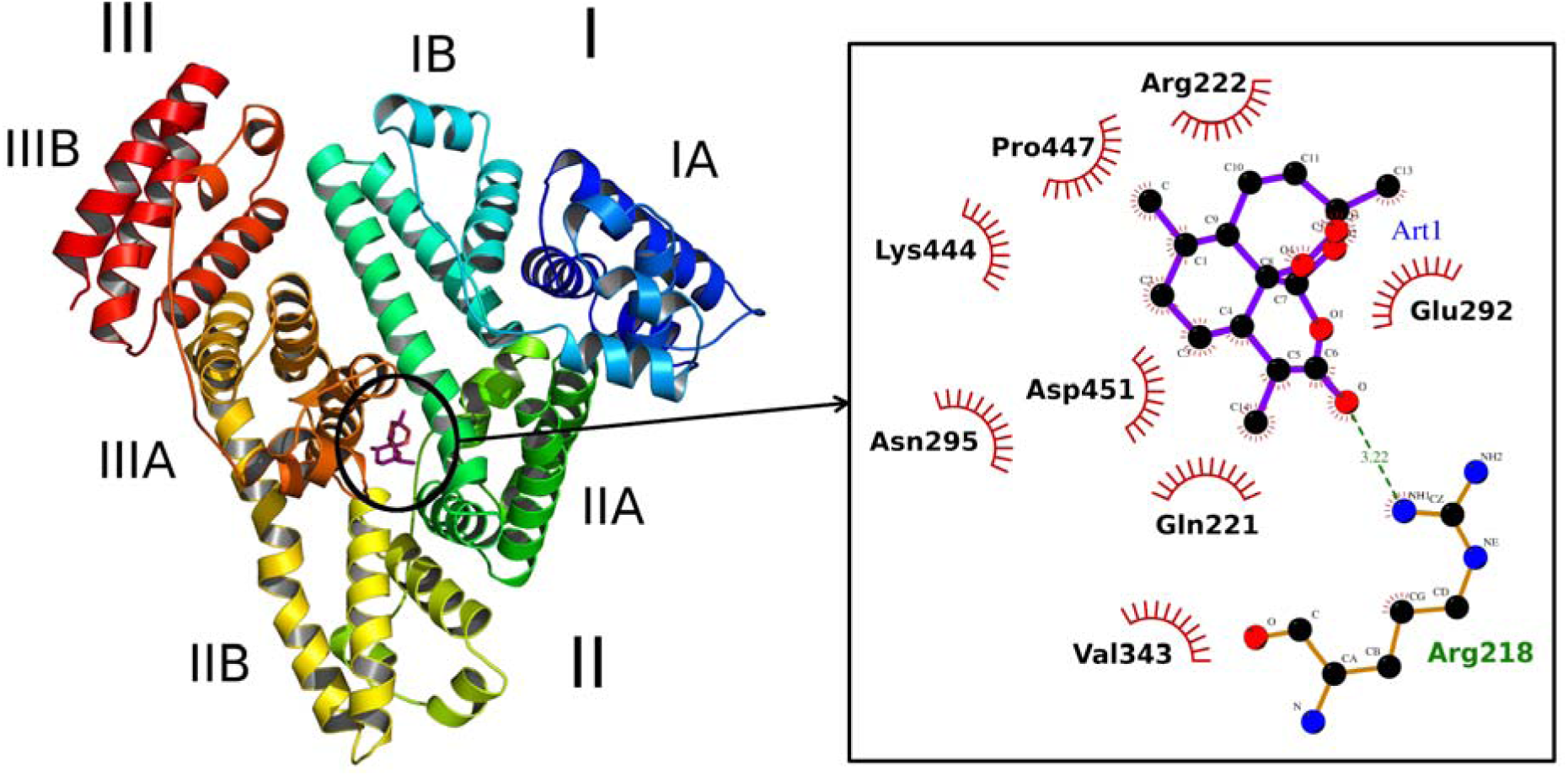
Docking of ART with the centroid conformation of the first cluster of HSA (left). Analysis of hydrophobic interactions and hydrogen bonds ART with HSA (right).

Since we consider ART as a new ligand for hGR, a comparative analysis of the interaction of artemisinin and dexamethasone with HSA was necessary. As in the case with ART, we carried out multiple docking simulations, PCA, cluster analysis and finally local docking of DEXA with the centroid conformation of HSA. In this case as well, docking simulations were launched 10 times with number of binding modes set to 20 for all 4 programs. PC decomposition was performed on the first two components using the COM coordinates (Fig. 7a). Later, cluster analysis was performed using the DBSCAN algorithm [49]. In the case of DEXA, for the DBSCAN parameters: minimum number of points was set to 5, and epsilon value was selected 4.0 (Fig. S4). As a result of the cluster analysis of DEXA with HSA, 10 clusters were obtained (Fig. S5). Only 1 cluster coincided in all 4 docking programs and is equal to 68.8% of all the DEXA conformations. This site coincides with the artemisinin binding site (Fig. 8a) and corresponds to the IIA domain (Drug site 1), which corresponds to experimental data [60].

**Fig. 7.**
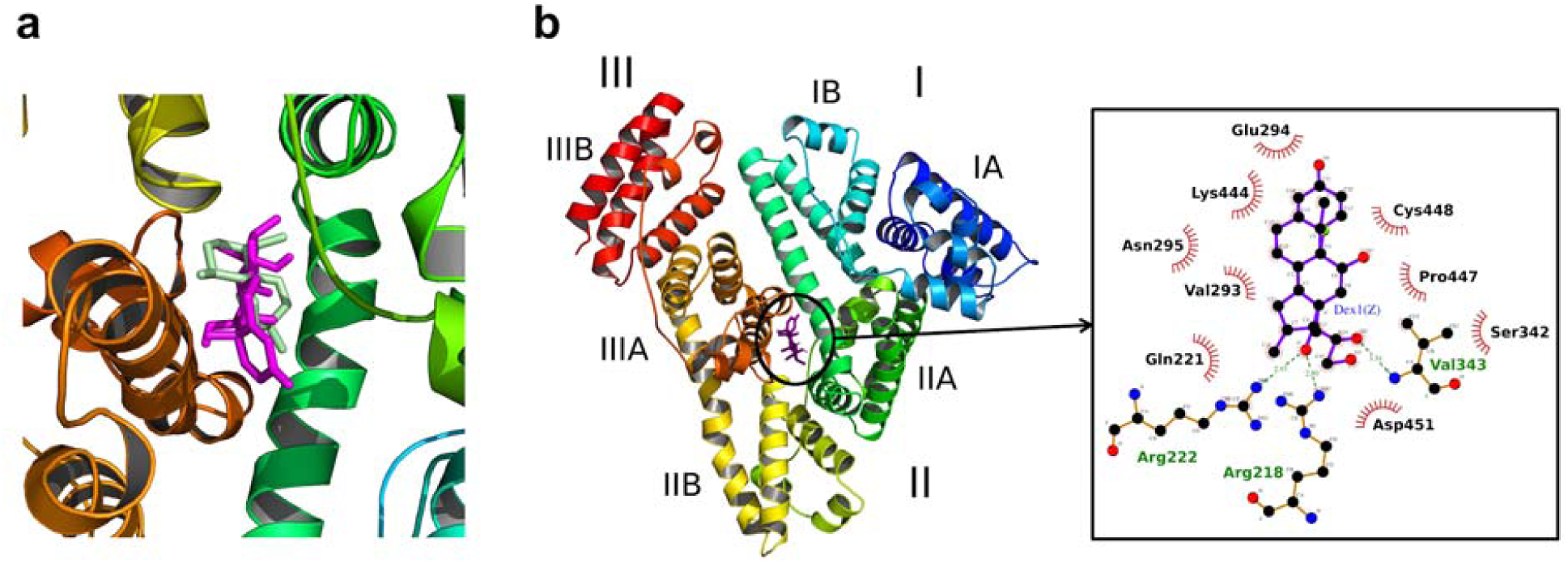
a) Local docking of HSA with ART (green) and DEXA (magenta). b) Local docking of DEXA with the centroid conformations of HSA (left). Analysis of hydrophobic interactions and hydrogen bonds of DEXA with HSA (right).

**Fig. 8.**
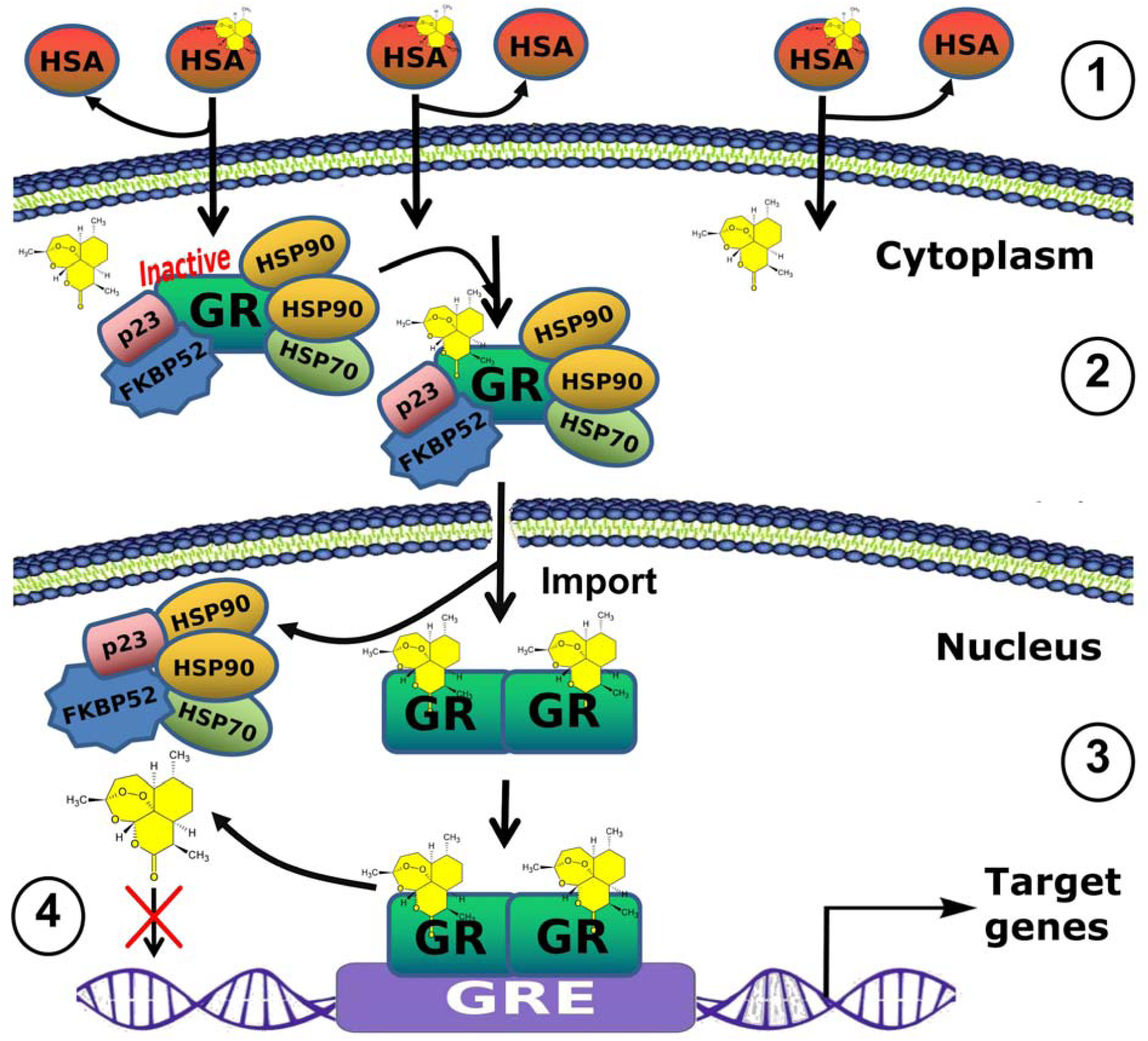
The hypothetical scheme: 1 - the interaction of ART with HSA and diffusion through the membrane; 2 - ART interaction with GR bonded with Hsp90 in cytoplasm; 3 - the penetration of complex through the nuclear pore, dimerization, the interaction with GRE on DNA with the expression of target genes; 4 - the absence of direct interaction with DNA.

Then, local docking was performed with this site using Autodock Vina [44] (Fig. 8b). According to the docking results, DEXA has a binding affinity of -9.3 kcal/mol, which is higher than the binding affinity of ART. This energy value is explained by the fact that DEXA forms 3 hydrogen bonds with HSA and hydrophobic interactions (Fig. 7b). The obtained results are consistent with *in vitro* study performed by Naik et al. [60].

DEXA forms hydrogen bonds with Arg218, Arg222 and Val343 and many hydrophobic interactions with amino acids from the IIA (Gln221, Val293, Glu294, Asn295 and Ser342) and IIIA (Lys444, Pro447, Cys448 and Asp451) domains. It is known that arginine 222 together with arginine 257 interact with the lactone carbonyl of warfarin, thereby stabilizing and orienting it in Drug site I [69].

## 4. Conclusion

Our spectral studies showed a decrease in the absorption peak of HSA at λ = 280 nm, which indicates an interaction between ART and HSA. Artemisinin also contributes to quenching of the fluorescence of HSA at λ = 337 nm, which correlates with its concentration. At the same time, ART does not fluoresce at maximum concentration. Therefore, such a decrease in intensity may be because of the quenching of the fluorescence of HSA by ART due to the formation of the complex. At the same time, there was only a slight shift of the fluorescence of HSA towards the short wavelengths, which indicates that the quenching was unchanged in the local dielectric medium of HSA due to tryptophan. Our data is consistent with the results of other study [25]. Within the studied concentration range, the results are consistent with the Stern-Volmer equation, as the graph shows a linear relationship. This suggests that in the presence of artemisinin, quenching of HSA proceeds according to a single mechanism. The quenching of the fluorescence of HSA with ART at different temperatures is, firstly, linear, and secondly, as the temperature rises, the extinction rate decreases, which indicates the static nature of extinguishing. To investigate the mechanism of interaction of ART with HSA, a docking analysis was performed.

Molecular docking was performed to clarify the binding modes of ART with HSA and the analyses of hydrophobic interactions and hydrogen bonds. It was shown that ART interacts with one sites of HSA, which is Drug site I. It forms hydrogen bonds with arginine at 218 position, while no hydrogen bonds are formed with tryptophan 214, which confirms the results of fluorescence. We have shown for the first time the formation of a hydrogen bond with arginine at 218 position, which plays a crucial role in the binding of drugs at site I, which includes thyroxine [67], warfarin, bilirubin and cholesterol [68]. Studies of the specificity of binding and modification of this arginine may be useful for therapeutic treatment methods that are aimed at preventing mediated adverse reactions in patients. [68]. We have shown that DEXA forms three hydrogen bonds (Arg218, Arg222 and Val343), while ART forms hydrogen bond with Arg218. It should be noted that many amino acids of HSA from IIA and IIIA subdomains that interact coincide for both ART and DEXA. The binding affinity of DEXA is higher than the binding affinity of ART.

In our previous study we carried out docking of ART with the monomer LBD of hGR, which showed that the main distribution of ART is on the interface formed by H3-H7-H10/H11 helices, which form the ligand binding pocket [16]. The interaction of a ligand with this sites leads to activation, for example, DEXA [18]. This site contains amino acids that are involved in the binding of GC, strong GR dimerization and play an important role in the recognitions of ligands and transactivation [18, 17].

Since GR can interact with ART, it is imported into the nucleus, it is extremely important to clarify that the results of our experiments show that there is no direct interaction of ART with the genomic DNA of S-180 cells. It was observed in concentrations ranging from 1-100 μM ART during incubation up to 48 hours.

Based on our data and in accordance with literary sources [4, 10, 17, 18, 25, 70], we propose a hypothetical scheme (Fig. 8).

So, based on the obtained results, we can assume that one of the main transporters of ART is HSA. Wherein, their interaction, binding sites, as well as energy parameters are similar to the corresponding parameters of dexamethasone. We have previously demonstrated that the interaction sites of ART with LBD of hGR coincide with those for DEXA [16]. This may represent the molecular basis for ligand-dependent activation of GR. In this case, the study of direct interaction of ART with genomic DNA is very important. We have shown the absence of direct interaction, thus ART could be considered as a new ligand (modulator, agonist) for GR. It can be assumed that the many activities of ART can be mediated by GR. Thus, our proposed scheme can explain the possible mechanism of action of ART, which includes the transportation through HSA and the biological effects - through GR.

## 5. Acknowledgements

Research was performed within the Government budget financing from Ministry of Education and Science of the Republic of Armenia, State Committee of Science (10-2-1-4). The research is carried out using the equipment of the shared research facilities of HPC computing resources at Lomonosov Moscow State University. Authors are thankful to the Laboratory of Toxinology and Molecular Systematics of L. A. Orbeli Institute of Physiology NAS RA.

