## Supplementary Materials for "*In vitro* and *in silico* identification of the mechanism of interaction of antimalarial drug – artemisinin with human serum albumin and genomic DNA"

**Appendix A. Supplementary Data**

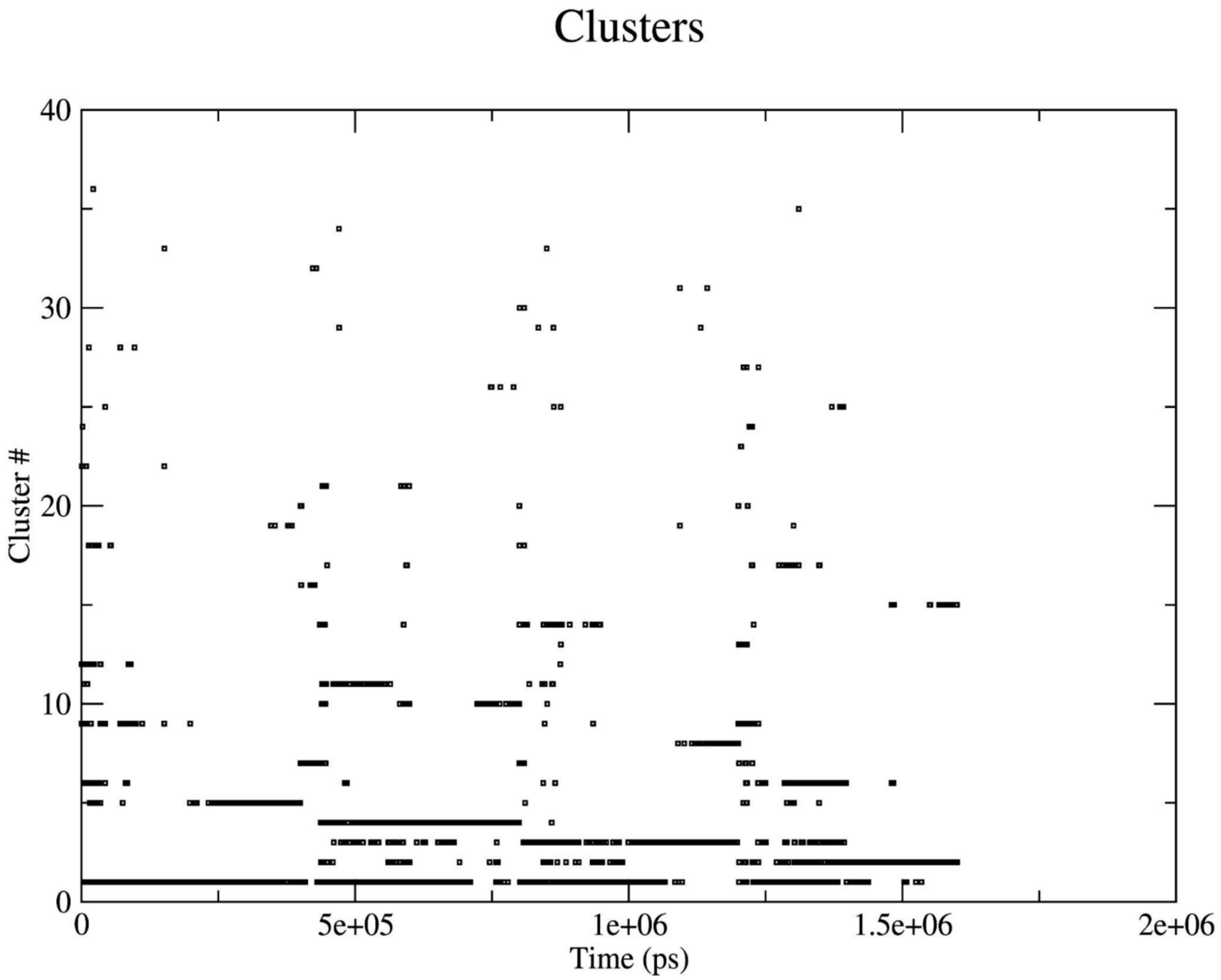


Fig. S1. Distribution of HSA conformations over time.

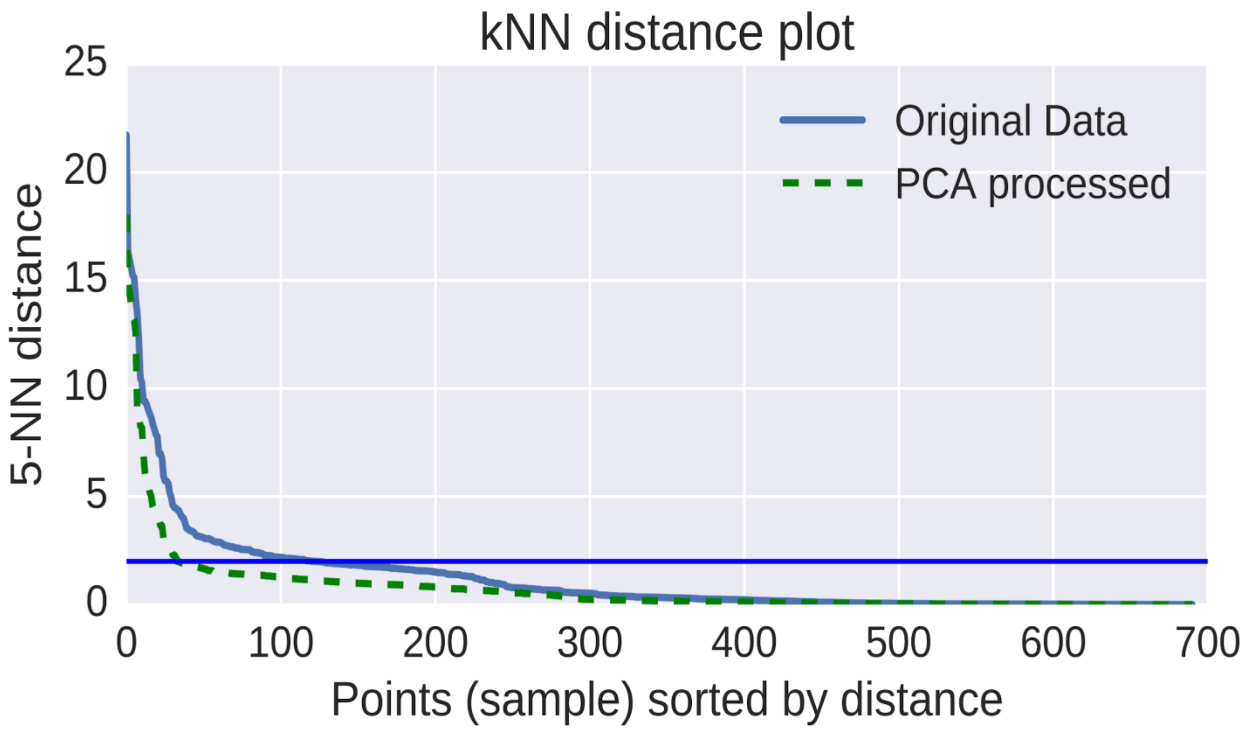


Fig. S2. k-Nearest Neighbor distance plot of the center of mass coordinates from the results of multiple docking runs of ART.

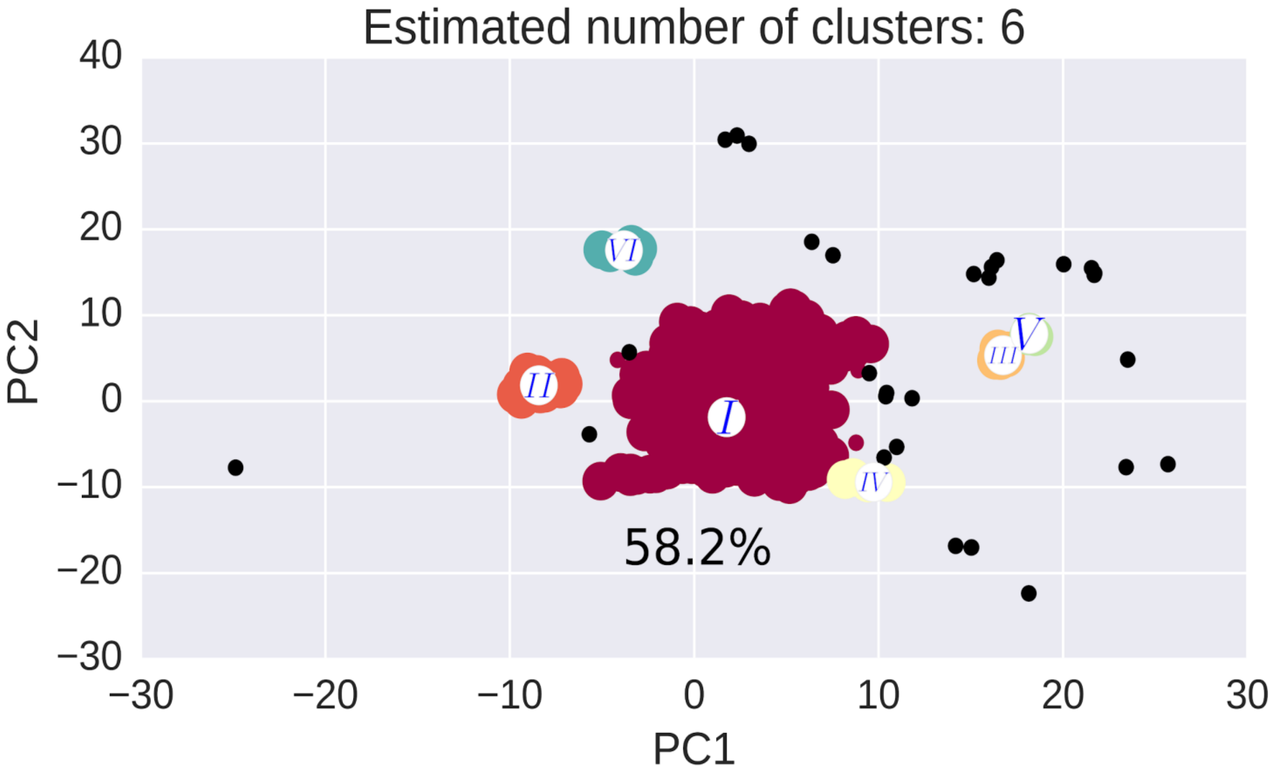


Fig. S3. Cluster analysis of the COM data from the docking results of ART with HSA using the DBSCAN algorithm. Roman numerals denote clusters, where the 1st cluster contains 58.2% of docking conformations.

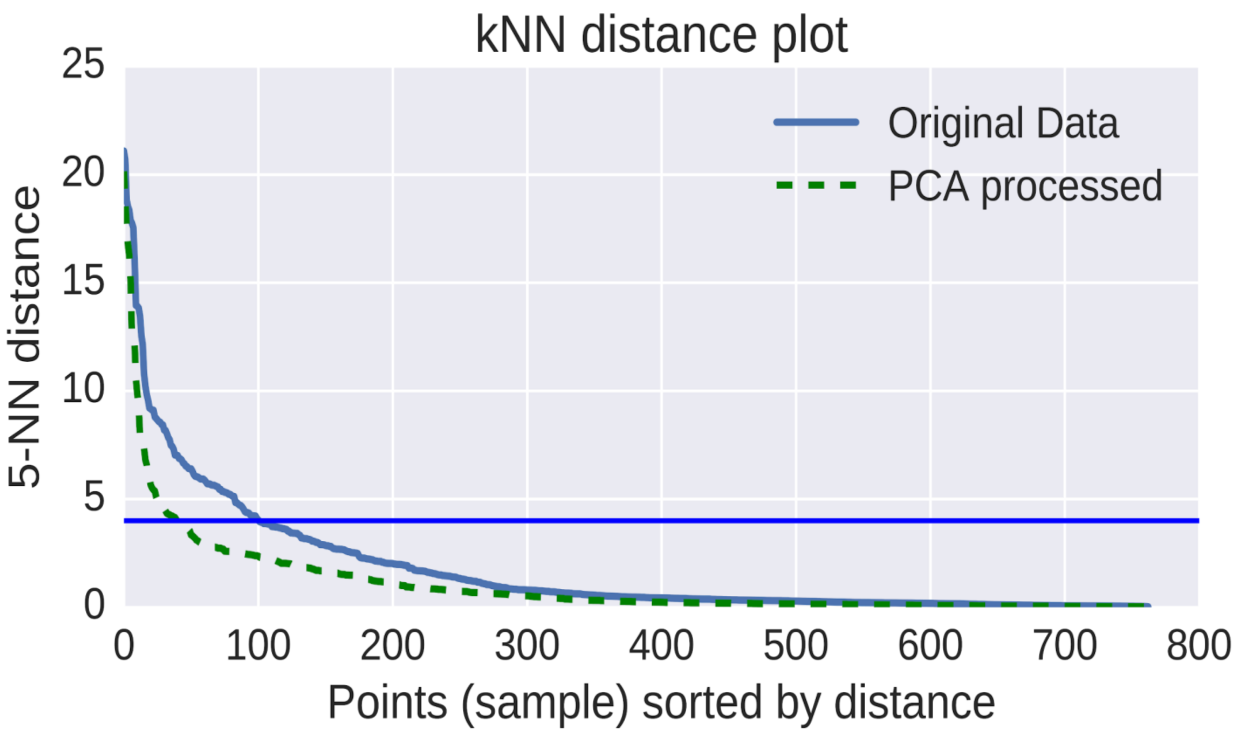


Fig. S4. k-Nearest Neighbor distance plot of the center of mass coordinates from the results of multiple docking runs of DEXA.

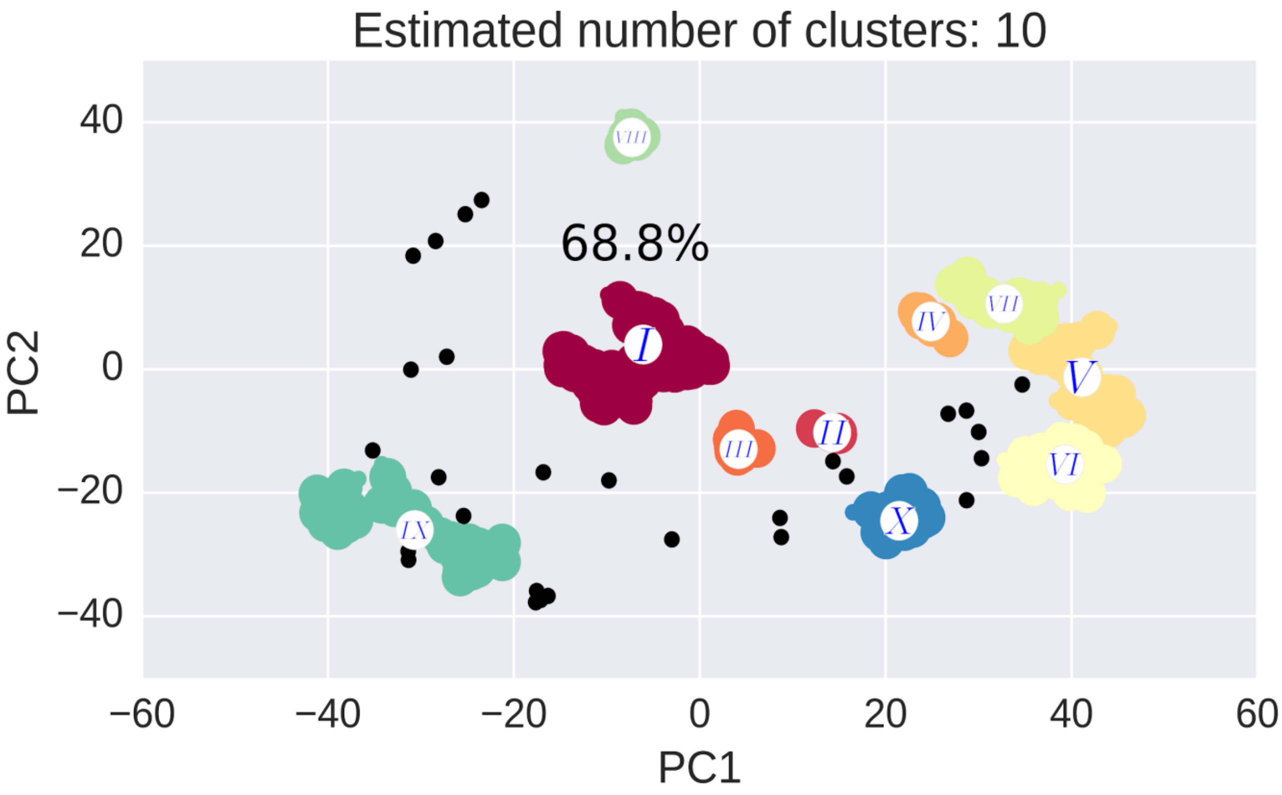


Fig. S5. Cluster analysis of the COM data from the docking results of DEXA with HSA using the DBSCAN algorithm. Roman numerals denote clusters, where the 1st cluster contains 68.8% of docking conformations.
